# Interpreting HIV Diagnostic Histories into Infection Time Estimates: Analytical Framework and Online Tool

**DOI:** 10.1101/323808

**Authors:** Eduard Grebe, Shelley N. Facente, Jeremy Bingham, Christopher D. Pilcher, Andrew Powrie, Jarryd Gerber, Gareth Priede, Trust Chibawara, Michael P. Busch, Gary Murphy, Reshma Kassanjee, Alex Welte, on behalf of the Consortium for the Evaluation and Performance of HIV Incidence Assays (CEPHIA)

## Abstract

**Background:** It is frequently of epidemiological and/or clinical interest to estimate the date of HIV infection or time-since-infection of individuals. Yet, for over 15 years, the only widely-referenced infection dating algorithm that utilises diagnostic testing data to estimate time-since-infection has been the ‘Fiebig staging’ system. This defines a number of stages of early HIV infection through various standard combinations of contemporaneous discordant diagnostic results, using tests of different sensitivity.

**Objective:** To develop a new, more nuanced infection dating algorithm, we generalised the Fiebig approach to accommodate positive and negative diagnostic results generated on the same *or* different dates, and arbitrary current or future tests – as long as the test sensitivity is known. For this purpose, test sensitivity is conceptualised as the probability that a specimen will produce a positive result, expressed as a function of time since infection. This can be summarised as a median ‘diagnostic delay’ parameter, together with a measure of inter-subject variability.

**Methods:** The present work outlines the analytical framework for infection date estimation using subject-level diagnostic testing histories, and data on test sensitivity. We introduce a publicly-available online HIV infection dating tool that implements this estimation method, bringing together 1) curatorship of HIV test performance data, and 2) infection date estimation functionality, to calculate plausible intervals within which infection likely became detectable for each individual. The midpoints of these intervals are interpreted as infection time ‘point estimates’ and referred to as Estimated Dates of Detectable Infection (EDDIs).

**Results:** In many settings, including most research studies, detailed diagnostic testing data are routinely recorded, and can provide reasonably precise estimates of the timing of HIV infection. We present a simple logic to the interpretation of ‘diagnostic testing histories’ into ‘infection time estimates’, either as a point estimate (EDDI) or an interval (earliest plausible to latest plausible dates of detectable infection), along with a publicly-accessible online tool that supports wide application of this logic.

**Conclusions:** This tool, available at https://tools.incidence-estimation.org/idt/, is readily updatable as test technology evolves, given the simple architecture of the system and its nature as an open source project.

## Background

For pathogenesis studies, diagnostic biomarker evaluation, and surveillance purposes, it is frequently of interest to estimate the HIV infection time of study subjects (i.e., the date of infection or time-since-infection). Ideally, a biomarker signature would provide reasonable direct estimates of an individual’s time-since-infection, but natural inter-subject variability of pathogenesis and disease progression makes this difficult. This work presents a general schema for utilising qualitative (i.e. positive/negative) diagnostic test results to estimate the time of HIV infection. Such estimates can be further refined by interpreting quantitative results on diagnostic or staging assays.(1)

Most simply, nuanced infection dating applies to subjects who produce a negative test result and also (usually at a later time) a positive test result, taking into account that no test can detect infection immediately after infectious exposure. Hence, infection can at best be estimated to have occurred during an interval in the past, relative to the date(s) of the test(s).

When a subject obtains discordant results, i.e. a negative and a positive test result on the same day, this typically manifests as positive results on ‘more sensitive’ tests than those on which the negative results were obtained. What we mean here by higher sensitivity is a shorter delay between acquisition and detectability of infection. For high-performing diagnostic tests, such as are normal for HIV and other viral infections like hepatitis C, test sensitivity is best understood as the probability of identifying a positive case as a function of time since infection (which is conventionally summarised as merely the probability of correctly identifying a positive case).

For more than 15 years, the only widely-referenced infection dating algorithm using diagnostic test results to estimate time-since-infection has been the ‘Fiebig staging’ system (2). This system defines a number of stages of early HIV infection through various standard combinations of contemporaneous discordant results using diagnostic tests of different sensitivity. For example, Fiebig stage 1 is defined as exhibiting reactivity on a viral load assay, but not (yet) on a p24 antigen assay, and in the seminal 2003 paper was estimated to begin approximately 11 days after infection, with a mean duration of 5.0 days (2). The particular tests used in these original calculations are largely no longer in use, nor commercially available. Others have used newer diagnostic assays to recalibrate the Fiebig stage mean duration estimates or define similar stages as an analogue to the Fiebig method (3, 4), though as tests evolve and proliferate, it becomes infeasible to calibrate all permutations of test discordancy.

Building from the Fiebig staging concept, we developed a new, more nuanced infection dating algorithm to meet the needs of a substantial collaboration (the Consortium for the Evaluation and Performance of HIV Incidence Assays – CEPHIA) in support of the discovery, development and evaluation of biomarkers for recent infection (5–7). The primary CEPHIA activity has been to develop various case definitions for ‘recent HIV infection’, with intended applicability mainly to HIV incidence surveillance rather than individual-level staging, although the latter application has also been explored (6, 8). CEPHIA has been able to identify large numbers of well characterized specimens and provide consistent conditions in which to conduct laboratory evaluations of several candidate incidence assays (5, 7) A key challenge faced by CEPHIA was that specimens in the repository had been collected from numerous studies, each of which used different diagnostic algorithms. The results of the tests performed in each of these studies therefore capture different information about the timing of HIV acquisition or seroconversion. To meet this challenge we had to link specimens from thousands of study-patient interactions into a coherent and consistent infection dating scheme, which enabled interpretation of arbitrary diagnostic test results (as long as the performance of the tests and questions were known). This general approach was first described in (9), but substantially refined in the present work.

In order to align diagnostic testing information across multiple sources, one needs a common reference event in a patient history – ideally, the time of an exposure that leads to infection. When dealing with actual patient data, however, we explicitly confront the point that in reality we are usually constrained to estimate the time when a particular test (X) would have first detected the infection. We will call this the test(X)-specific *Date of Detectable Infection*, or DDI_X_.

The present work outlines the analytical framework for infection date estimation using ‘diagnostic testing histories’, and introduces a publicly-available online HIV infection dating tool that implements this estimation, bringing together 1) curatorship of HIV test performance data, and 2) infection date estimation functionality. It is readily updatable as test technology evolves, given the simple general architecture of the system and its nature as an open source project.

## Methods

### Generalised Fiebig-like staging

The fundamental feature of the Fiebig staging system (2) is that it identifies a naturally-occurring sequence of discordant diagnostic tests that together indicate early clinical disease progression. The approximate duration of infection can be deduced from the categorization into stages through analysis of the combination of specific assay results.

As we demonstrate below, it is most robust to interpret any combination of diagnostic test results into an estimated duration of infection, if these tests have been independently benchmarked for diagnostic sensitivity (i.e. a median or mean duration of time from infection to detectability on that assay has been estimated). This more nuanced method allows both for incorporation of results from any available test, and from results of tests run on specimens taken on different days.

Interpreted at the population level, a particular test’s sensitivity curve expresses the probability that a specimen obtained at some time *t* after infection will produce a positive result. The key features of a test’s sensitivity curve (represented by the purple curve in Figure 1) are that:

- there is effectively no chance of detecting an infection immediately after exposure;
- after some time, the test will almost certainly detect an infection;
- there is a characteristic time range over which this function transitions from close to zero to close to one. This can be summarised as something very much like a mean or median and a standard deviation.

**Figure 1.**
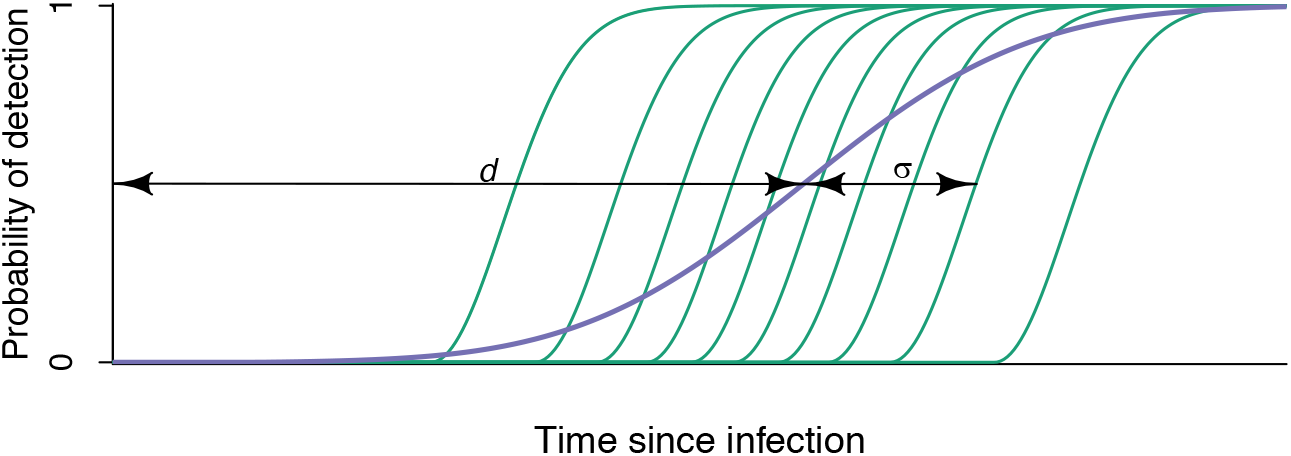

Various host and pathogen attributes, such as concurrent infections, age, the particular viral genotype, post-infection factors, etc., affect the performance of a test for a particular individual. This in principle determines a subject-specific curve, such as one of the green curves in Figure 1, which capture the probability, as a function of time, that specimens from a particular subject will produce a positive diagnostic result. Because assay results are themselves imperfectly reproducible even on the same individual, these green curves do not transition step-like from zero to one but have some finite window of time over which they transition from close to zero to close to one.

In contrast to the usual statistical definition of ‘sensitivity’ as the proportion of ‘true positive’ specimens that produce a positive result, we summarise the population-level sensitivity of any particular diagnostic test into one or two ‘diagnostic delay’ parameters (*d* and *σ* in Figure 1). By far the most important parameter is an estimate of ‘*median diagnostic delay*.’ In Figure 1, this is the parameter *d*. If there were perfect test result conversion for all subjects (i.e. no assay ‘noise’), and no inter-subject variability, this would reduce the smoothly varying purple curve to a step function.

To estimate individual infection times, then, one needs to obtain estimates of the median diagnostic delays for all tests occurring in a data set, and then interpret each individual assay result as excluding some ‘non-possible’ segment of time, ultimately resulting in a final inferred interval of time during which infection likely occurred. The prototypical situations in which one can perform dating, within this paradigm, are then when a subject:

1. tests positive at a given time after testing negative at some earlier time, or
2. tests positive on some component of an algorithm, and negative on another component, contemporaneously.

These calculations require that each individual has at least one negative test result and at least one positive test result. In the primitive case where there is precisely one of each, namely a negative result on a test with an expected diagnostic delay of *d*_1_ at *t*_1_ and a positive result on a test with an expected diagnostic delay of *d*_2_ at *t*_2_, then the interval is simply from (*t*_1_ − *d*_1_) to (*t*_2_ − *d*_2_). When there are multiple negative results on tests at 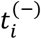 each with a diagnostic delay 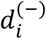, and/or multiple positive results on tests at 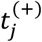 each with a diagnostic delay 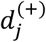, then each individual negative or positive test result provides a candidate earliest plausible and latest plausible date of infection. The most informative tests, then, are the ones that functionally narrow the ‘infection window’ (i.e. result in the latest start and earliest end of the window) by excluding periods of time during which infection would not plausibly have become detectable. In this case, the point of first ‘detectability’ refers to the time when the probability of infection being detected by an assay first exceeds 0.5.

These remaining plausible ‘infection windows’ are usually summarised as intervals, the midpoint of which is naturally considered a ‘point estimate’ of the date of infection. These intervals can be understood as plateaus on a very broadly plateaued (rather than ‘peaked’) likelihood function, as shown in Figure 2. Given a uniform prior, this can be interpreted as a Bayesian posterior, with [*a, b*] in Figure 2 showing the 95% credibility interval (i.e. the interval encompassing 95% of the posterior probability density). Such a posterior, derived from an individual’s diagnostic testing history, could also serve as a prior for further analysis, if there is an available quantitative biomarker for which there is a robustly calibrated maturation/growth curve model. We do not deal with this in the present work, but it is explored elsewhere (1), and is an important potential application of this framework and tool.

**Figure 2.**
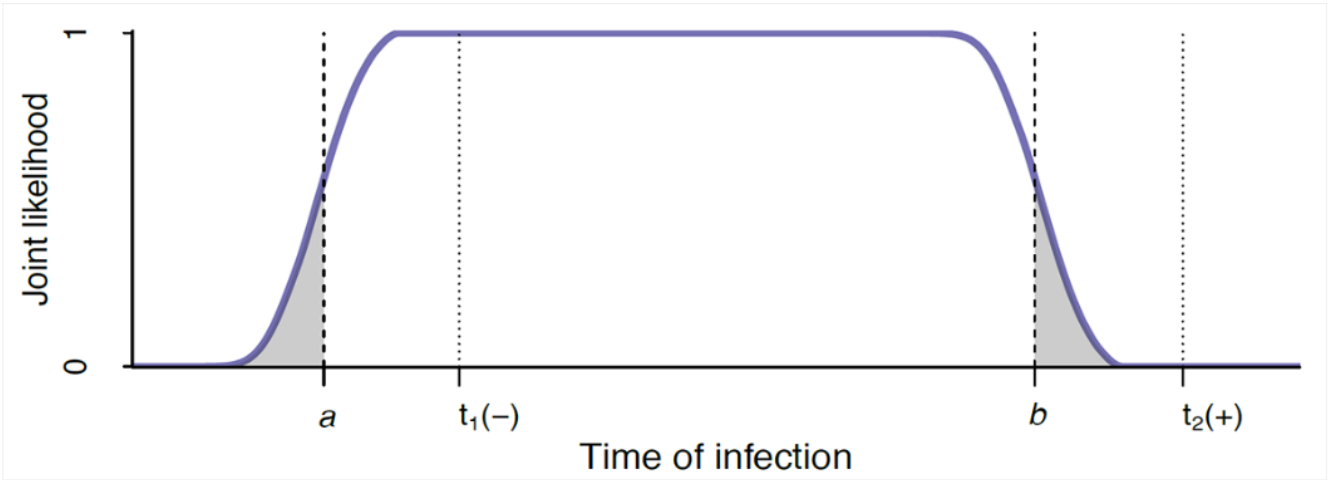

In the next section we derive a formal likelihood function – i.e. a formula capturing the probability of seeing a data element or set (in this case, the set of negative and positive test results), given hypothetical values of the parameter(s) of interest – here, the time of infection.

### Formal likelihood function and impact of test correlation

It is analytically useful to specify an explicit ‘likelihood function’, i.e. a formula for capturing the probability of seeing a data element (or set), given some hypothetical values for parameters which determine the behaviour of the underlying system, including the measurement process. This facilitates all the usual statistical manipulations for obtaining confidence intervals, Bayesian posteriors, etc. For the present application, test sensitivity curves such as those in Figure 1 are precisely the likelihood of obtaining a positive result upon application of a given test, at a given time since infection. The likelihood of obtaining a negative result, on this very application of the test, is simply 1 minus the likelihood of obtaining a positive result, i.e. a vertically flipped version of the test sensitivity curve. As noted above, meaningful infection dating relies on having at least one negative test result and at least one positive test result.

#### Classical test converson series

To begin, we consider precisely one negative and one positive test result, arising from two subject-study interactions, at times *t*_1_ and *t*_2_ respectively, separated by some duration *δ.* In order to make inferences about the time of infection, we construct a likelihood function which expresses the probability of seeing these two particular results, *as a function of a hypothetical infection time.* This kind of likelihood (of two observations) is usually written as the product of:

- the likelihood of seeing one result (chosen arbitrarily to be considered first) given the hypothetical time of infection, and
- the likelihood of seeing the other result, given

◦ the same hypothetical time of infection, and
◦ the fact that the other result has in fact been obtained.

Using *T_inf_* to denote the actual time of infection that we are trying to estimate, *t_inf_* to denote particular values of hypothetical infection time and [+, *t_n_*] and [−, *t_n_*] to denote positive and negative test results at observation times *t*_n_, respectively, this can be written as:

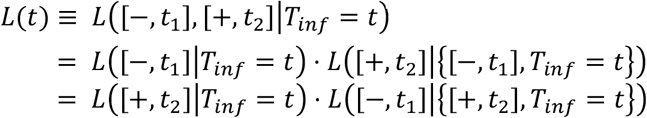

capturing that the likelihood of seeing both of two events (A and B) is equal to either

1. *the likelihood of A* multiplied by *the likelihood of B, given A*, i.e. *L*(*A*) · *L*(*B*|*A*), or
2. *the likelihood of B* multiplied by *the likelihood of A, given B*, i.e. *L*(*B*) · *L*(*A*|*B*).

The details of the *conditioned* likelihoods, which might be complex, must necessarily be such that the two formulations are equivalent. We will focus in detail on the first formulation, as it seems more intuitively appealing when *t*_1_ < *t*_2_.

Figure 3 shows, in thick green and red, respectively, the *population-level* likelihoods of observing the negative test result at *t*_1_ and the positive test result at *t*_2_. A subset of the family of *individual-level* curves, chosen to visually suggest their distribution, is indicated as thin lines. A close look at these curves reveals that they are the horizontally flipped (and in the case of the green curves, also vertically flipped) test sensitivity curves of the tests performed (compare with the detailed shapes in Figure 1). These curves display information for each test result, considered independently.

**Figure 3.**
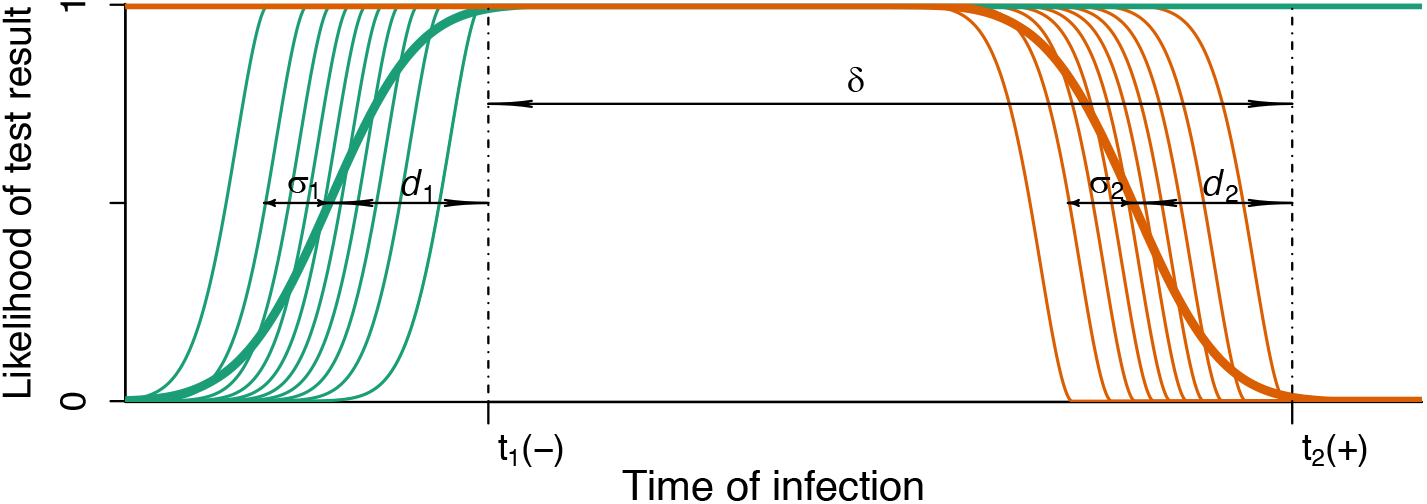

The fundamental point of estimating an infection time is that both tests were in fact performed on the same individual. It is highly likely that those individuals who convert rapidly, post infection, on Test_1_ also convert rapidly on Test_2_ – which might, after all, be the same test, and is likely to be a similar test. The details of this conditioning can in principle be complex, and it is infeasible to study all the correlations between all tests in use in studies. A critical question, then, is whether, when, and how this correlation impacts the conditioned likelihoods which are the fundamental building block of a forma inference of infection time from diagnostic testing histories.

The ‘worst case’ scenario would be when the correlation is very strong, as it would be if the tests performed at the two times are in fact the same test. We have explicitly implemented a model of test sensitivity based on the following points:

- the performance of any test is defined by a family of *N* individual-level sensitivity curves of the type in Figures 1 and 3.
- for a particular test, each individual-level curve is a shifted Weibull with the same shape and scale parameter.
- The shift parameter is normally distributed, though with a discretised realisation, with step 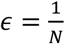 – i.e. we assign individual diagnostic delays (Weibull shift parameters) to the percentiles 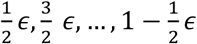 of a normal distribution.
- The mean of the distribution of shift parameters is a test’s mean diagnostic delay (*d* in Figures 1 and 3), and the standard deviation (*σ* in Figures 1 and 3) manifests as something akin to a shape parameter of the population-level curve.

To keep the scenario simple initially, we first consider the case when the two test times differ by more than *d* + *σ*. We later consider the complication of the other extreme, i.e. when the positive and negative test results are obtained on the same day, and the distributions of the diagnostic delays overlap substantially.

The behaviour of the fully-conditioned likelihood expression

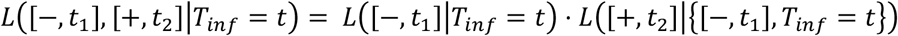

can then be understood by considering how the factor *L*([+, *t*_2_]|{[−, *t*_1_], *T_inf_* = *t*}) might differ from the naïve population-averaged, unconditioned *L*([+, *t*_2_]|*T_inf_* = *t*). The latter is what one can obtain from a study investigating the performance of one or several diagnostic tests, without having to apply the particular test combinations to particular individuals.

We now analyse the various ranges of *t_inf_* which are qualitatively different from each other:

##### Values of t_inf_ on the far-left end of the timeline

For very ‘early’ hypothetical infection times, the likelihood of seeing the negative result at *t*_1_ becomes very small. If that negative result has indeed occurred at *t*_1_, it would normally be the result of a laboratory error, which would have a reasonable chance of being detected with strong quality controls. If the error remains undetected, testing positive at *t*_2_ (a time later than *t*_1_ by a significant margin) is nevertheless almost assured, i.e.
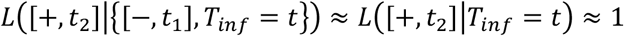

So, in this case there is no discernible difference between the unconditioned and conditioned likelihood, although there is no analytical cure for a false negative result.

##### Values of t_inf_ within the dynamic range of the Test_1_ sensitivity curve

Figures 4a-4e consider a series of hypothetical infection times (*t_inf_*) that span the likely range of diagnostic delays.

**Figure 4.**
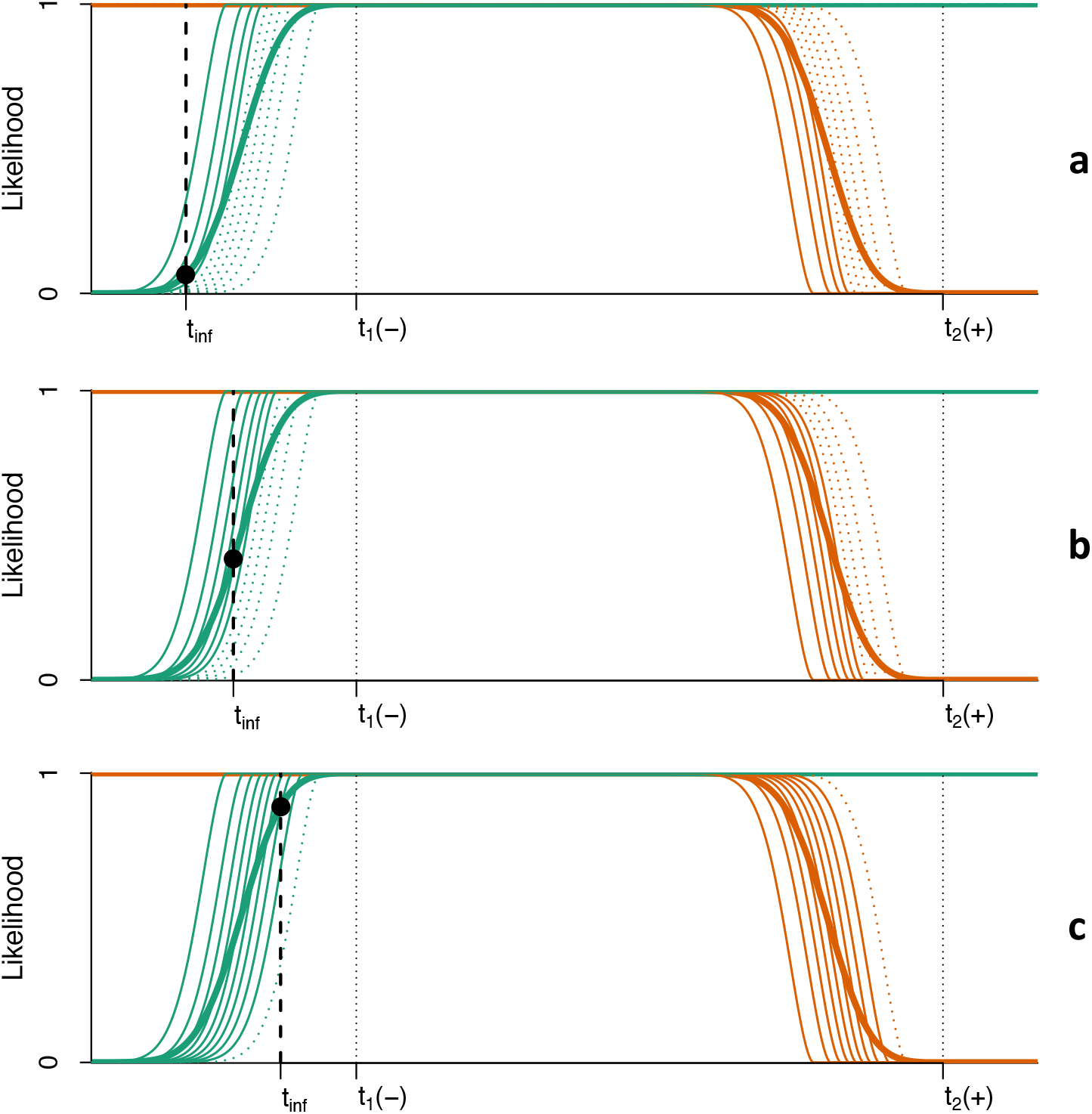
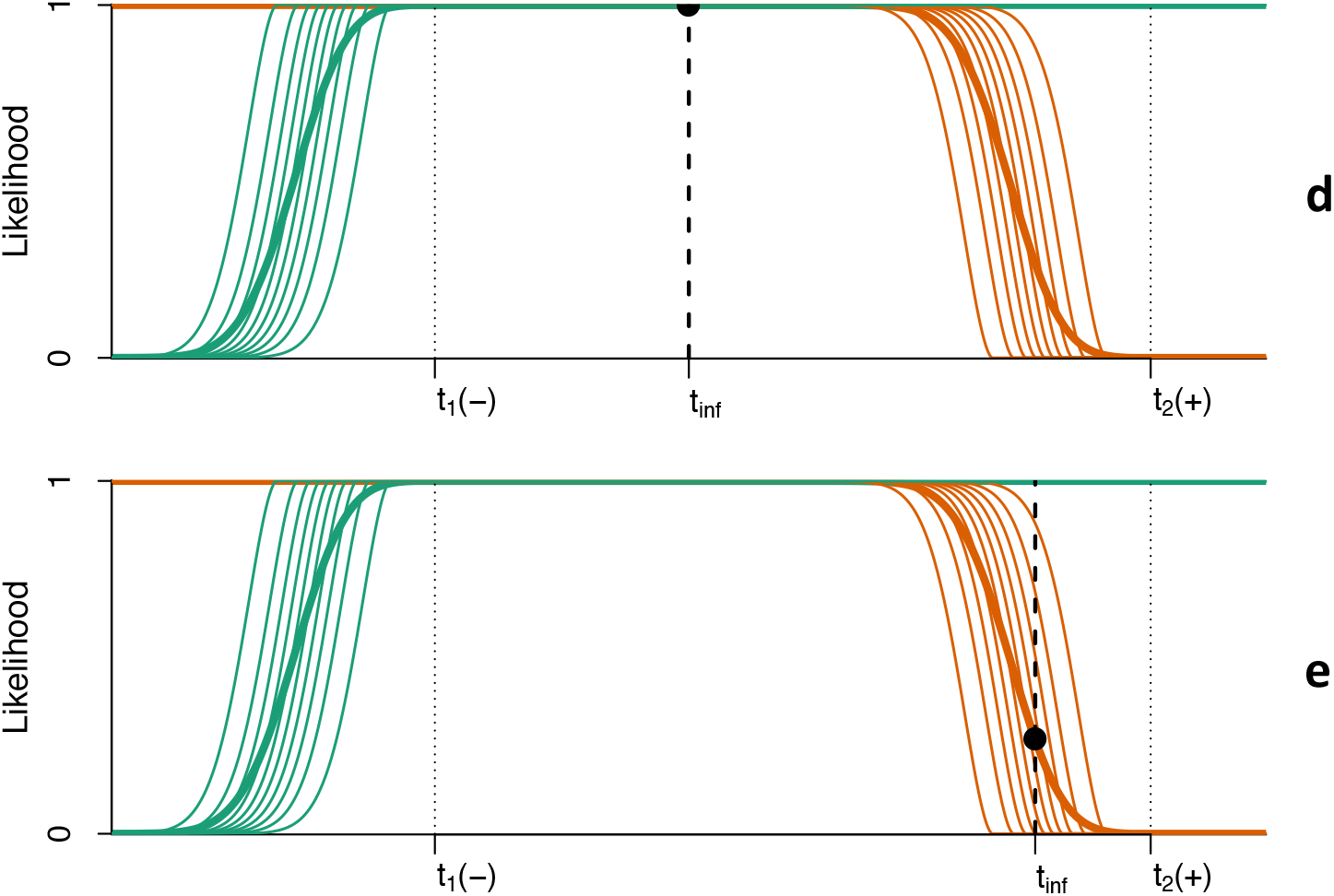

Figure 4a, indicating that the negative result at *t*_1_ occurs somewhat longer after infection than the mean diagnostic delay, is suggestive of the subject being a significantly slower-than-average progressor on the diagnostic marker. This is captured by the dotted green (faster) individual progression curves for Test_1_, indicating their reduced plausibility. Correspondingly, only the slowest progression rates are plausible among the red curves for Test_2_. Nevertheless, given the location of *t_inf_*, namely long before the application of Test_2_, it does not matter which of the Test_2_ progression curves the individual is likely to be on – they all evaluate to 1 so long after infection.

Figure 4b, indicating that the negative result at *t*_1_ occurs at a time after infection approximately equal to the mean diagnostic delay, is suggestive of the subject not being a significantly faster-than-average progressor on the diagnostic marker. This is captured by the reduced number of dotted green (fastest) individual progression curves for Test_1_. Correspondingly, the fastest progression rates are less plausible among the red curves for Test_2_. Once more, given the location of the hypothetical infection time, namely long before the application of Test_2_, it does not matter which of the Test_2_ progression curves the individual is likely to be on – they all evaluate to 1 so long after infection.

Figure 4c, indicating that the negative result at *t*_1_ occurs at a time after infection that is significantly less than the mean diagnostic delay, is consistent with all but one of the green and hence red individual progression lines are plausible. Not only does the negative result at *t*_1_ not imply significant conditioning on the subject’s diagnostic marker progression rate, but all the individual-level red curves in any case evaluate to 1 *at the time of* Test_2_.

##### Values of t_inf_ anywhere near, or to the right of t_1_

For these ‘later’ hypothetical infection times, we *expect* to see a negative result for the test at *t*_1_, even more so than in Figure 4c, and so, the negative result provides no information on the question of whether the subject is prone to rapid or slow test conversion. Hence, no modification is implied of *L*([+, *t*_2_]|{[−, *t*_1_], *T_inf_* = *t*}) relative to the population average *L*([+, *t*_2_]|*T_inf_* = *t*), though of course in this region there are many values of *t_inf_* for which this likelihood is not approximately 1. Figures 4d and 4e show values of *t_inf_* on the ‘plateau’ and on the ‘descent’ from the plateau in the dynamic range of diagnostic delays of Test_2_.

The three zones of *t_inf_* discussed above account for the full range of values of *t* for which the joint likelihood is to be constructed. It is clear that the full joint likelihood is indeed given by the product of the unconditioned population-level likelihoods for the two test results, as shown in Figure 5. As the curves obtain values indistinguishable from either 0 or 1 for much of their range, this product is little more than a superposition of the two curves.

**Figure 5.**
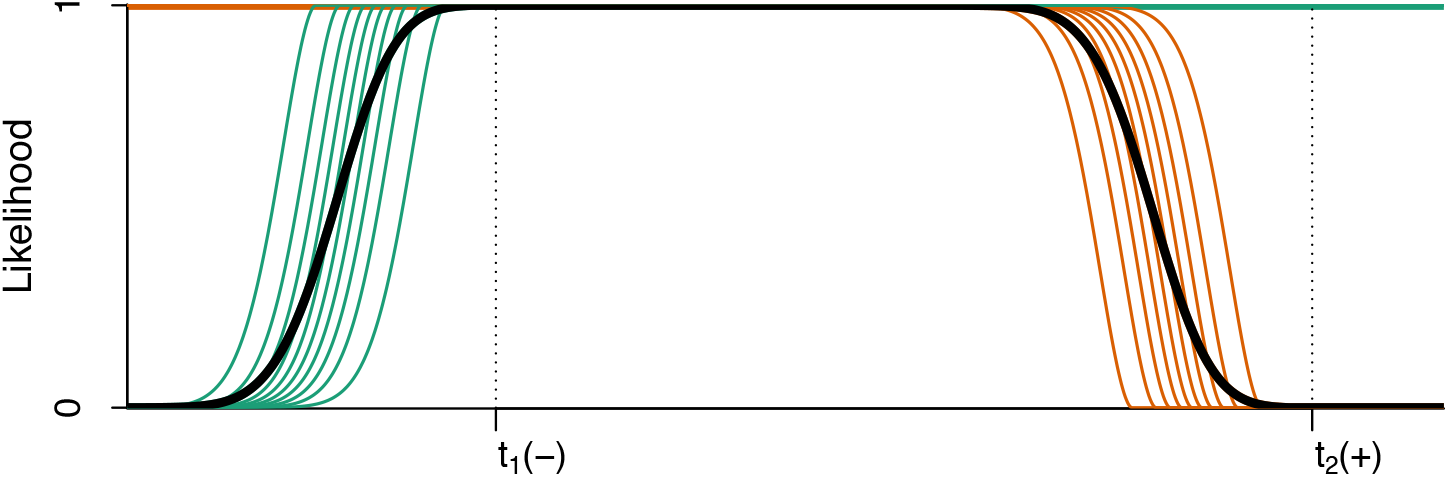

Figure 2, shown earlier, indicates where this round-shouldered plateau is located relative to the test dates and population-averaged diagnostic delays. Under the parametric assumptions outlined above, one may specify a ‘confidence level’ (such as usually encapsulated in a significance level *α*, chosen to be 0.05 in Figure 2) and calculate the bounds of the (in our case, 95%) ‘credibility interval’ [*a, b*], encompassing the relevant proportion of the posterior probability density *p*(*t*). We therefore find the values of *a* and *b* that satisfy

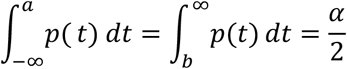

Note that when *t*_1_(−) and *t*_1_(+) are separated by a substantial period of time, the credibility interval is likely to be narrower than naïve bounds defined simply by the population-average diagnostic delays. When the period is short, the credibility bounds are likely to shift outward from the naïve bounds.

#### Discordant results on a given study-visit

Figure 6 shows the typical ‘discordant test’ situation, where a test with a longer diagnostic delay produces a negative result and a test with a shorter diagnostic delay produces a positive result, at the same visit.

**Figure 6.**
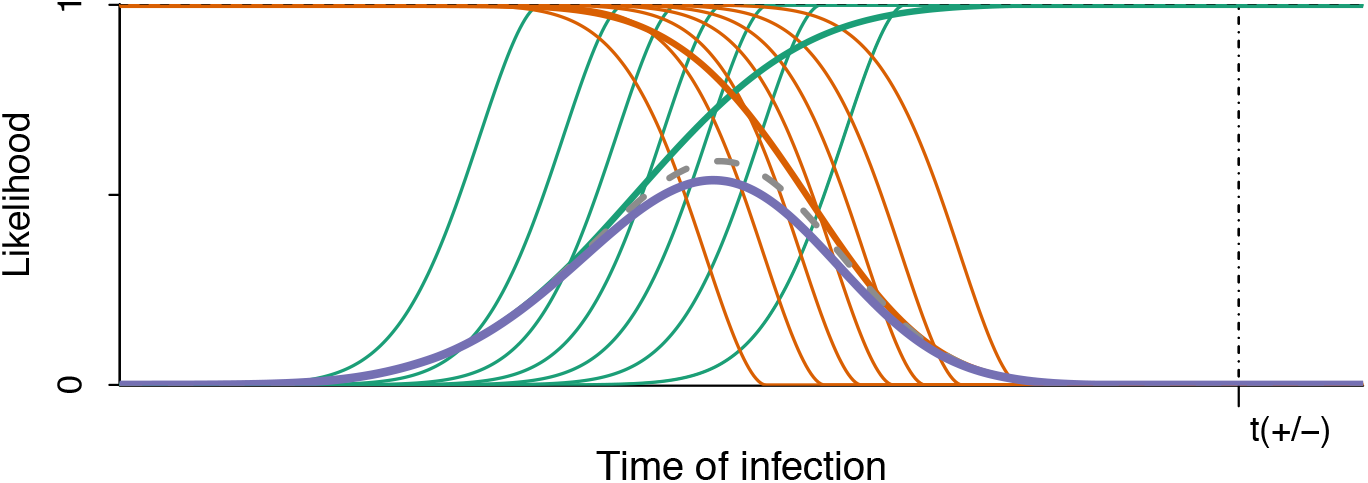

Even here, though not as starkly as in the case where the two tests are conducted at significantly different times, conditioning one result on the other has relatively modest impact. Moving the hypothetical infection time to the left, the negative result becomes less likely, and the effect of the conditioning on the likelihood of seeing the second test result becomes more significant. However, as the hypothetical infection time moves further left, the times under consideration leave the dynamic range of the positive test, and it becomes ever less plausible that a negative test result is obtained. We do not explicitly display figures indicating the conditioning implied for various hypothetical values of infection time, but merely indicate in the solid blue curve the formally calculated fully specified joint likelihood which takes this conditioning into account in terms of the extreme correlation model outlined above. This exact likelihood does not differ meaningfully from the simple product of the population-level likelihoods of the two tests (shown dashed, in grey). The main conclusion, then, is that relative to the test date, plausible infection times are largely located between the two diagnostic delays (with some spreading due to variability).

Figure 7 shows the situation where the dynamic ranges of the tests are essentially the same. In this case, the plausible dates of infection are centred around the shared diagnostic delay of the tests, again with some spread for variability. The relatively small amplitude of the exact curve indicates that the fully conditioned discordancy is significantly less likely to occur than one would infer from a naïve calculation, but the key point is that the infection time estimate is not affected at the level at which it can be plausibly reported.

**Figure 7.**
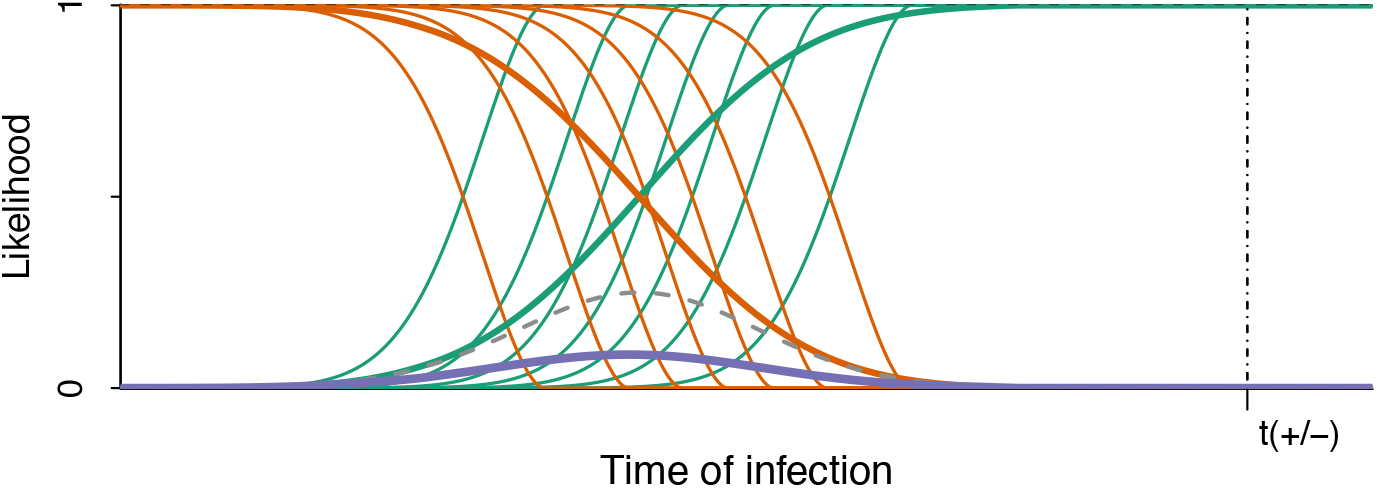

Figure 8 shows an outlier situation in which a more sensitive test is negative while a less sensitive test is positive. Relative to the naïve product of likelihoods, the correctly specified joint likelihood is very small for all values of *t*. This indicates that such anomalous discordant results are extremely rare, arising most plausibly from test error. If such an outlier occurs without test error, the fully-conditioned likelihood could differ significantly from the naïve one; however, it would depend on essentially unknowable details of distributional tails and test correlation, and such cases are sufficiently rare to have no impact on conclusions drawn from observing large numbers of individuals. Note that the extreme rarity of anomalous discordant results is a function of the very strong intra-test correlation assumed in this model; in reality the intra-test correlation is likely far less strong, making these events less rare but also lessening the discrepancy between the naïve and fully-conditioned likelihood.

**Figure 8.**
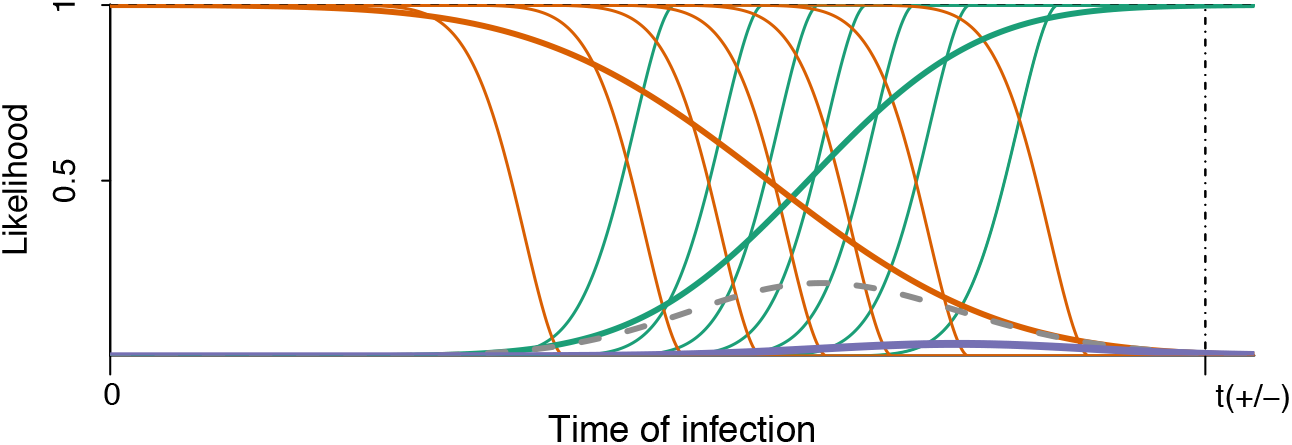

## Implementation

The public online Infection Dating Tool is available at https://tools.incidence-estimation.org/idt/. The source code for the tool is available publicly under the GNU General Public License Version 3 open source licence at doi:10.5281/zenodo.1488117.

In practice, the timing of infectious exposure is seldom known, even in intensive studies, and studies of diagnostic test performance therefore provide *relative* times of test conversion (10–12). Diagnostic delay estimates are therefore anchored to a standard reference event – the first time that a highly-sensitive viral load assay with a detection threshold of 1 RNA copy/mL of plasma – would detect an infection. We call this the *Date of Detectable Infection* (DDI). The tool endows study subjects with a point estimate of this date, which we call the *Estimated Date of Detectable Infection* (EDDI). The time from infectious exposure to DDI is likely to be variable between individuals, but the tool does not rely on any assumptions about the average duration of this pre-DDI state. Details and evaluation of the performance of the diagnostic delay estimates underlying this tool compared with other methods for estimation of infection dates are available elsewhere (1).

The key features of our online tool for HIV infection date estimation are that:

1. Users access the tool through a website where they can register and maintain a profile which saves their work, making future calculations more efficient.
2. Individual test dates and positive/negative results, i.e. individual-level ‘testing histories’, not just algorithm-level diagnoses, can be uploaded in a single comma-delimited text file for a group of study subjects.
3. Estimates of the relative ‘diagnostic delay’ between the assays used and the reference viral load assay must be provided, with the option of using a curated database of test properties which provides cited estimates for over 60 HIV assays.
a. If a viral load assay’s detection threshold is known, this can be converted into a diagnostic delay estimate via the exponential growth curve model (1, 2). We assume that after the viral load reaches 1 RNA copy/mL, viral load increases exponentially during the initial ramp-up phase. The growth rate has been estimated at 0.35 log_10_ RNA copies/mL per day (i.e., a doubling time of slightly less than one day) (2). The growth rate parameter defaults to this value, but users can supply an alternative estimate.
4. Using the date arithmetic described above, when there is at least one negative test result and at least one positive test result for a subject, the uploaded diagnostic history results in:

a. a point estimate for the date of first detectability of infection (the EDDI);
b. an earliest plausible and latest plausible date of detectable infection (EP-DDI and LP-DDI); and
c. the number of days between the EP-DDI and LP-DDI (i.e., the size of the ‘DDI interval’), which gives the user a sense of the precision of the estimate.

### Access / User profiles

Anyone can register as a user of the tool. The tool saves users’ data files as well as their choices about which diagnostic delay estimates to use for each assay, both of which are only accessible to the user who uploaded them. No person-identifying information is used or stored within the tool; hence, unless the subject identifiers being used to link diagnostic results can themselves be linked to people (which should be ruled out by pre-processing before upload) there is no sensitive information being stored on the system.

### Uploading diagnostic testing histories

A single data file would be expected to contain a ‘batch’ of multiple subjects’ diagnostic testing histories. Conceptually, this is a table like the fictitious example in Table 1, which records that:

- one subject (Subject A) was seen on 10 January 2017, at which point he had a detectable vial load on an unspecified qualitative viral load assay, but a negative Bio-Rad Geenius™ HIV-1/2 Supplemental Assay (Geenius) result
- another subject (Subject B) was screened negative using a point-of-care (PoC) rapid test (RT) on 13 September 2016, and then, on 4 February 2017, was confirmed positive by Geenius, having also tested positive that day on the PoC RT

**Table 1.**

| **Subject** | **Date** | **Test** | **Result** |
| --- | --- | --- | --- |
| Subject A | 2017-01-10 | Qualitative VL | Positive |
| Subject A | 2017-01-10 | Geenius | Negative |
| Subject B | 2016-09-13 | POC RT | Negative |
| Subject B | 2017-02-04 | POC RT | Positive |
| Subject B | 2017-02-04 | Geenius | Positive |

In order to facilitate automated processing, the tool demands a list of column names as the first row in any input file. While extraneous columns are allowed without producing an error, there must be columns named *Subject, Date, Test* and *Result* (not case sensitive). Data in the subject column is expected to be an arbitrary string that uniquely identifies each subject. Dates must be in the standard ISO format (YYYY-MM-DD).

It is fundamental to the simplicity of the algorithm that assay results be either ‘positive’ or ‘negative’. There are a small number of tests, notably Western blot and the Geenius, which sometimes produce ‘indeterminate’ results (partially, but not fully, developed band pattern). Note that there is some lack of standardisation on interpretation of the Western blot, with practice differing in the United States and Europe, for example. While we provide default values for common Western blot assays, users may enter appropriate estimates for the specific products and interpretations in use in their specific context.

We now briefly reconsider Table 1 by adding the minor twist that the Geenius on Subject B is reported as indeterminate. In this case, the data must be recorded as results on either one or both of two separate tests:

1. a ‘Test-Indeterminate’ version of the test – which notes whether a subject will be classified either as negative, or ‘at least’ as indeterminate; and
2. a ‘Test-Full’ version of the test, which determines whether a subject is fully positive or not.

There is then no longer any use for an un-suffixed version of the original test. The data from Table 1 is repeated in Table 2 with differences highlighted. The only changes are the use of the Test-Indeterminate version for Subject A’s negative Geenius result and an indeterminate Geenius result for Subject B. Note that even while Subject A’s test results have not changed, their testing history now looks different, as completely negative results are reported as being negative even for the condition of being indeterminate. Subject B’s indeterminate result on 4 February requires two rows to record, one to report that the test result is not fully negative (positive on ‘Geenius Indeterminate’), and one to report that the result is not fully positive (negative on ‘Geenius Full’). Once diagnostic delays are provided for these two sub-tests, the calculation of infection dates can proceed without any further data manipulation on the part of the user.

**Table 2.**
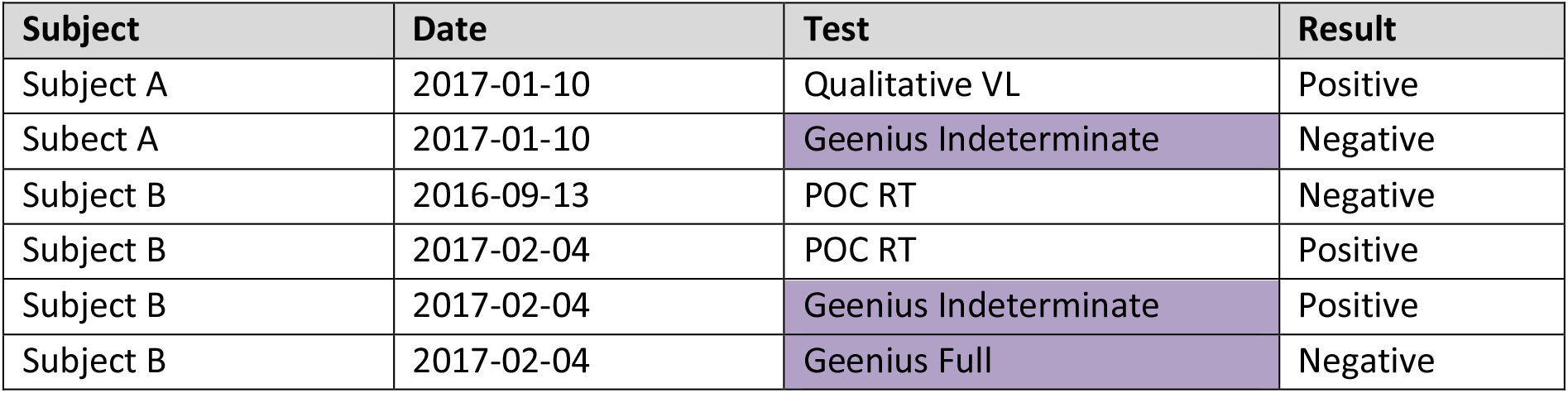

| Subject | Date | Test | Result |
| --- | --- | --- | --- |
| Subject A | 2017-01-10 | Qualitative VL | Positive |
| Subect A | 2017-01-10 | Geenius Indeterminate | Negative |
| Subject B | 2016-09-13 | POC RT | Negative |
| Subject B | 2017-02-04 | POC RT | Positive |
| Subject B | 2017-02-04 | Geenius Indeterminate | Positive |
| Subject B | 2017-02-04 | Geenius Full | Negative |

### Provision of test diagnostic delay estimates

As described above, tests are summarised by their diagnostic delays. The database supports multiple diagnostic delay estimates for any test, acknowledging that these estimates may be provisional and/or disputed. The basic details identifying a test (i.e. name, test type) are recorded in a ‘tests’ table, and the diagnostic delay estimates are entered as records in a ‘test-properties’ table, which then naturally allows multiple estimates by allowing multiple rows which ‘link’ to a single entry in the tests table. A test property entry captures the critical parameter of the ‘average’ (usually median) diagnostic delay obtained from experimental data and, when available, a measure of the variability of the diagnostic delay (denoted *σ*).

The system’s user interface always ensures that for each user profile, there is exactly one test property estimate, chosen by the user, as ‘in use’ for infection dating calculations at any point in time. Users need to ‘map’ the codes occurring in their data files (i.e. the strings in the ‘Test’ column of uploaded data files) to the tests and diagnostic delay estimates in the database, with the option of adding entirely new tests to the database, which will only be visible to the user who uploaded them. The tool developers welcome additional test estimates submitted for inclusion in the system-default tests/estimates.

### Execution of infection dating estimation

The command button ‘process’ becomes available when an uploaded testing history has no unmapped test codes. Pressing the button leads to values, per subject, for EP-DDI, LP-DDI, EDDI, and DDI interval, which can be previewed on-screen and downloaded as a comma-delimited file.

By default, the system employs simply the ‘average’ diagnostic delay parameter, in effect placing the EP-DDI and LP-DDI bounds on the DDI interval where the underlying sensitivity curve evaluates to a probability of detection of 0.5. When the size of the inter-test interval (*δ*) is greater than about 20 times the diagnostic delay standard deviation (*σ*), this encompasses more than 95% of the posterior probability.

As an additional option, when values for both *d* and *σ* are available, and under the parametric assumptions outlined earlier, users may specify a significance level (*α*), and the system will calculate the bounds of a corresponding credibility interval. The bounds of the central 95% (in the case of *α* = 0.05) of the posterior are labelled the EP-DDI and LP-DDI.

### Database Schema

This tool makes use of a relational database, which records information in a set of linked tables, including:

- **subjects**: This table captures each unique study subject, and after infection date estimation has been performed, the subject’s EDDI, EP-DDI, LP-DDI and DDI interval size.
- **diagnostic_test_history**: This table records each test performed, by linking to the subjects table and recording a date, a ‘test code’, and a result. During the estimation procedure, a field containing an ‘adjusted date’ is populated, which records the candidate EP-DDI (in the case of a negative result) or LP-DDI (in the case of a positive result) after the relevant diagnostic delay has been applied to the actual test date.
- **diagnostic_tests**: This is a lookup table listing all known tests applicable to the current purposes (both system-provided and user-provided).
- **test_property_estimates**: This table records diagnostic delay estimates (system and user-provided). It allows estimates per test, with system default estimates flagged.
- **test_property_mapping**: This table records user-specific mapping of test codes by linking each test code in the diagnostic_test_history table to a test in the diagnostic_tests table, as well as the specific test property estimate ‘in use’ by that user for the test in question.

A number of subsidiary tables also exist to manage users of the system and allow linking of personal data files, maps, tests, and test property estimates to specific users.

## Results

### Example of infection date estimates from testing history data

A hypothetical example showing source data and the resulting infection date estimates is provided below. The example data are available with the source code (in a file named ExampleData.csv). Table 3A shows the testing history data file, which lists all diagnostic test results obtained for three subjects, which represent typical cases: Subject A had discordant test results on a single date, with the more sensitive test producing a positive result and the less sensitive test a negative result. Subject B seroconverted between two dates separated by some months. Subject C had a large number of tests, and first produced negative results, then discordant results (positive only on a NAT assay), then an immature antibody response, and finally exhibited a fully reactive Western Blot. A time series of this kind provides a detailed view of early disease stage progression and yields very precise infection time estimates.

**Table 3A:**
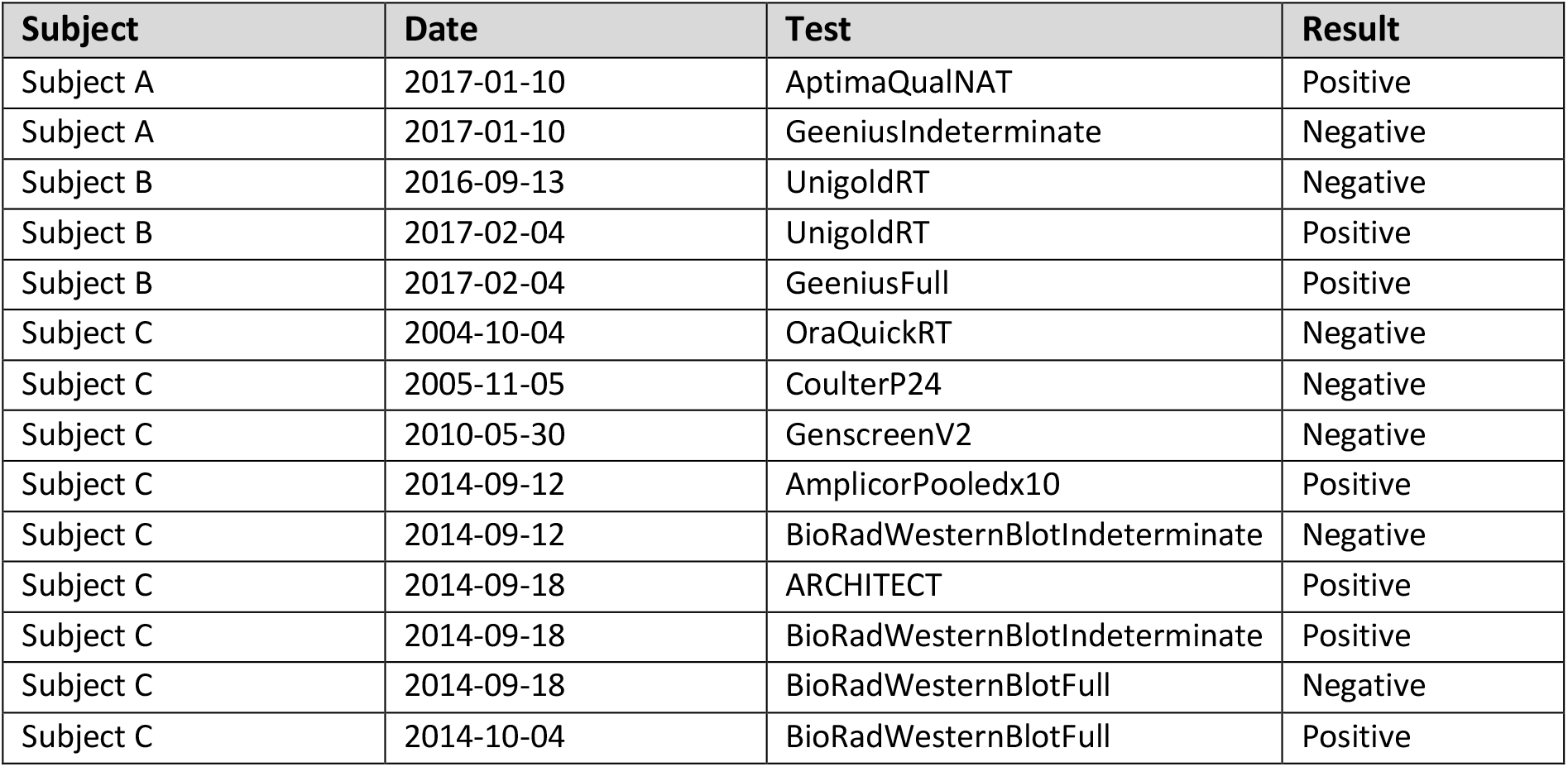
Example Dataset

| Subject | Date | Test | Result |
| --- | --- | --- | --- |
| Subject A | 2017-01-10 | AptimaQualNAT | Positive |
| Subject A | 2017-01-10 | GeeniusIndeterminate | Negative |
| Subject B | 2016-09-13 | UnigoldRT | Negative |
| Subject B | 2017-02-04 | UnigoldRT | Positive |
| Subject B | 2017-02-04 | GeeniusFull | Positive |
| Subject C | 2004-10-04 | OraQuickRT | Negative |
| Subject C | 2005-11-05 | CoulterP24 | Negative |
| Subject C | 2010-05-30 | GenscreenV2 | Negative |
| Subject C | 2014-09-12 | AmplicorPooledx10 | Positive |
| Subject C | 2014-09-12 | BioRadWesternBlotIndeterminate | Negative |
| Subject C | 2014-09-18 | ARCHITECT | Positive |
| Subject C | 2014-09-18 | BioRadWesternBlotIndeterminate | Positive |
| Subject C | 2014-09-18 | BioRadWesternBlotFull | Negative |
| Subject C | 2014-10-04 | BioRadWesternBlotFull | Positive |

Table 3B shows the mapping of test codes to tests in the tool’s database, together with median diagnostic delay estimates provided as default estimates in the database.

**Table 3B:**
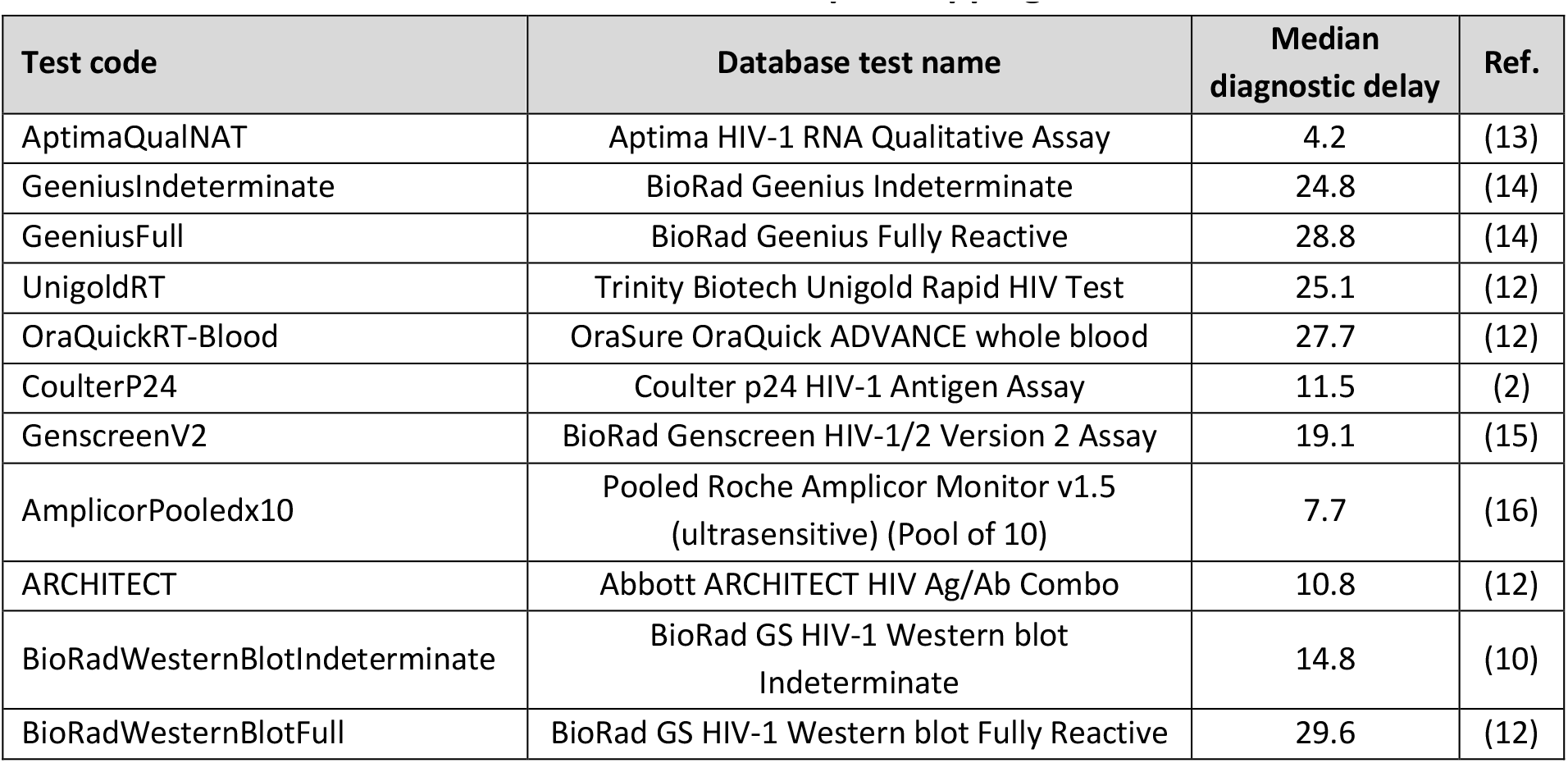
Example Mapping

| Test code | Database test name | Median diagnostic delay | Ref. |
| --- | --- | --- | --- |
| AptimaQualNAT | Aptima HIV-1 RNA Qualitative Assay | 4.2 | (13) |
| GeeniusIndeterminate | BioRad Geenius Indeterminate | 24.8 | (14) |
| GeeniusFull | BioRad Geenius Fully Reactive | 28.8 | (14) |
| UnigoldRT | Trinity Biotech Unigold Rapid HIV Test | 25.1 | (12) |
| OraQuickRT-Blood | OraSure OraQuick ADVANCE whole blood | 27.7 | (12) |
| CoulterP24 | Coulter p24 HIV-1 Antigen Assay | 11.5 | (2) |
| GenscreenV2 | BioRad Genscreen HIV-1/2 Version 2 Assay | 19.1 | (15) |
| AmplicorPooledx10 | Pooled Roche Amplicor Monitor v1.5 (ultrasensitive) (Pool of 10) | 7.7 | (16) |
| ARCHITECT | Abbott ARCHITECT HIV Ag/Ab Combo | 10.8 | (12) |
| BioRadWesternBlotIndeterminate | BioRad GS HIV-1 Western blot Indeterminate | 14.8 | (10) |
| BioRadWesternBlotFull | BioRad GS HIV-1 Western blot Fully Reactive | 29.6 | (12) |

Table 3C shows the results of the estimation procedure, together with a column indicating which test results were most informative for deriving the EP-DDIs and LP-DDIs.

**Table 3C:** Example Results

| Subject | EP-DDI<br>(naïve) | LP-DDI<br>(naïve) | Interval<br>size<br>(naïve) | EP-DDI<br>(95% CI) | LP-DDI<br>(95% CI) | EDDI<br>(95% CI<br>midpoint) | Interval<br>size<br>(95% CI) | *Most informative tests* |
| --- | --- | --- | --- | --- | --- | --- | --- | --- |
| Subject A | 2016-12-16 | 2017-01-06 | 21 | 2016-12-11 | 2017-01-05 | 2016-12-23 | 25 | *GeeniusIndeterminate_Neg 2017-01-10*<br>*AptimaQualNAT_Pos 2017-01-10* |
| Subject B | 2016-08-19 | 2017-01-06 | 140 | 2016-08-21 | 2017-01-03 | 2016-10-27 | 135 | *UnigoldRT_Neg 2016-09-13*<br>*GeeniusFull_Pos 2017-02-04* |
| Subject C | 2014-08-28 | 2014-09-04 | 7 | 2014-08-24 | 2014-09-05 | 2014-08-30 | 12 | *BioRadWesternBlot-Indeterminate_Neg 2014-09-12*<br>*BioRadWesternBlotFull_Pos 2014-10-04* |

Note that the most informative tests are those that exclude the greatest periods of time preceding (in the case of a negative result) and the period following (in the case of a positive result) the earliest dates of plausible detectability, calculated from the test’s diagnostic delay. These are not necessarily the tests performed on the last date on which a negative, or the first date on which a positive result was obtained.

Further note that when the testing interval is small, the 95% credibility interval tends to be wider than the naïve median-based DDI interval (Subjects A and C in the example), but when the testing interval is large, the credibility interval tends to be narrower than the naïve DDI interval (Subject B in the example).

### Use of the tool in real-world research studies

The infection dating tool described in this work has been utilized to estimate infection dates for all subjects who contributed specimens to the CEPHIA repository, where diagnostic testing histories could be obtained. A key aspect of that consortium’s work has been to characterise tests for recent HIV infection (HIV ‘incidence assays’) – in particular by estimating the two critical performance characteristics, the mean duration of recent infection (MDRI) and false-recent rate (FRR), which would not have been possible without individual-level infection time estimates, for example (8).

### Source code, distribution and modification

The whole code base for the tool is available in a public source code repository (at https://github.com/SACEMA/infection-dating-tool/, with the latest release always available at doi:10.5281/zenodo.1488117), and so anyone can deploy their own copy of the tool, or ‘fork’ the repository (i.e. make their own copy of the code repository) and make any modifications they wish. The only condition is that the origin of the code is acknowledged, and dissemination of the modified code is also in open source form under the same licensing. The developers of the tool welcome contributions to the code, which can be proposed through ‘pull requests’ issued on the source code hosting platform. Test characteristics for more than 60 common HIV diagnostic tests are included in the code base and are easy to update as new data become available.

Consistent infection dating could be of interest in the study of other infections. Only minor modifications and a database of tests and test property estimates would be required to deploy a separate version of the system handle other infections. This would be especially useful in contexts where multiple diagnostic platforms or algorithms have been used within a single dataset intended for a unified analysis.

## Conclusions

Consistent dating of infection events across subjects has obvious utility when analysing multi-site datasets that contain different underlying screening algorithms. Consistent use of ‘diagnostic history’ information is also valuable for individual-level interpretation of infection staging at diagnosis.

Even in intensive studies from which ‘diagnostic delay’ estimates are drawn, it is rarely possible to determine the actual date of infectious exposure. We have adopted a nomenclature based on the earliest date on which an infection would have had 50% probability of being detected, using a viral load assay with a detection threshold of 1 copy per ml, and we refer to this as the Date of Detectable Infection (DDI).

A limitation of this approach is that it relies on details of diagnostic testing histories that are often not recorded or clearly reported. For example, it may be noted that a subject produced a negative Western blot result on a particular date, but without recording of the specific product and the interpretive criteria employed. This challenge is further compounded by country-specific variations in assay names and interpretive criteria for the same assays. Self-reported testing histories may also lack precise information on the dates of tests and the specific assays employed, in which case this tool cannot be used to estimate infection time. When a last negative result and a first positive result are separated by a long period of time (e.g. two years), very uninformative infection time estimates are produced. In these cases, the interpretation of additional quantitative markers – utilising the infection time intervals estimated by this tool as ‘priors’ – can yield informative estimates (1).

A simple method for interpreting additional quantitative markers (such as a signal-to-cutoff ratio from the ARCHITECT diagnostic assay or a normalised optical density from the Limiting Antigen Avidity recency assay) would be to interpret the obtained result using a Mean Duration of Recent Infection vs. recency discrimination threshold calibration curve to derive a ‘time scale’– i.e. on average, a subject producing *y* quantitative result has been infected for less than *x* days, see for example (8).

However, in many settings, including most research studies, detailed diagnostic testing data are routinely recorded, and especially when regular testing occurred, can provide reasonably precise estimates of the timing of HIV infection even with purely qualitative results.

We have presented a simple logic to the interpretation of ‘diagnostic testing histories’ into ‘infection time estimates’, either as a point estimate (EDDI) or an interval (EP-DDI – LP-DDI), along with a publicly-accessible online tool that supports wide application of this logic.

## List of Abbreviations

CEPHIA –: Consortium for the Evaluation and Performance of HIV Incidence Assays
CI –: Credibility Interval
DDI –: Date of Detectable Infection
EDDI –: Estimated Date of Detectable Infection
EP-DDI –: Earliest Plausible Date of Detectible Infection
HIV –: Human Immunodeficiency Virus
LP-DDI –: Latest Plausible Date of Detectable Infection
PoC –: Point-of-Care
RNA –: Ribonucleic Acid
RT –: Rapid Test

## Declarations

### Availability of materials and data

The example dataset analysed during this study is published in its full form in this article and also available with the source code of the tool. All source code is available from a public repository under an open source license, using the persistent DOI: https://doi.org/10.5281/zenodo.1488117.

Other datasets analysed using this tool, including CEPHIA data, are not publicly available, since they contain personally identifying information, notably actual dates of HIV test results. Anonymised data with modified dates can be obtained from the corresponding author upon reasonable request.

### Author contributions

AW: First draft of manuscript, overall project leadership; AW, EG and SNF: conceptualisation, data curatorship, software design, code development, writing of the manuscript; JB: Visualisation; RK, MPB, GM, CDP: conceptualisation; AP, JG, GP, TC: code development. AP was the primary software developer.

## Acknowledgements

The authors would like to thank Kevin P. Delaney (US Centers for Disease Control and Prevention) for insightful comments on the draft manuscript and assistance with obtaining test property estimates.

## Appendix A Web interface layout overview

Once logged in, the system presents users with four primary pages, accessible via links spread in horizontal tabs below the header, as shown in Figure A.1. The first three are described in turn below, with the fourth the subject of a separate publication (Welte, et al., forthcoming).

**Figure A.1:**
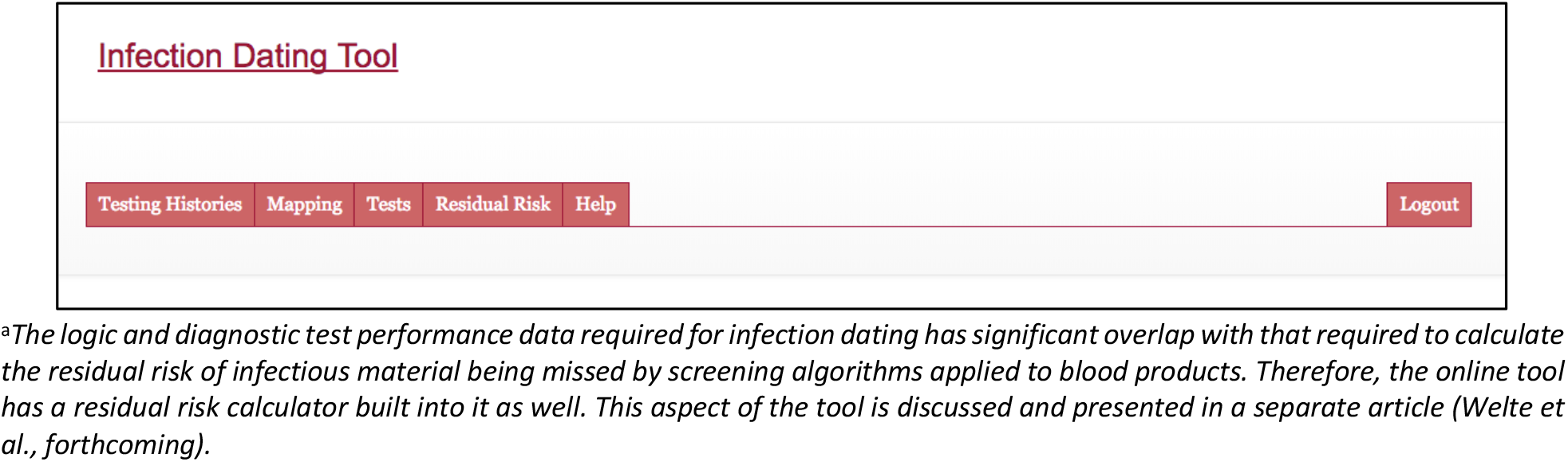
Navigation^a^

### Testing Histories

This tab (Figure A.2) allows users to locate, view and delete previously uploaded ‘testing histories’, and to upload new ones. It is also where users trigger the action of processing the uploaded testing histories into ‘infection dating estimates’, which can then be viewed and downloaded.

**Figure A.2:**
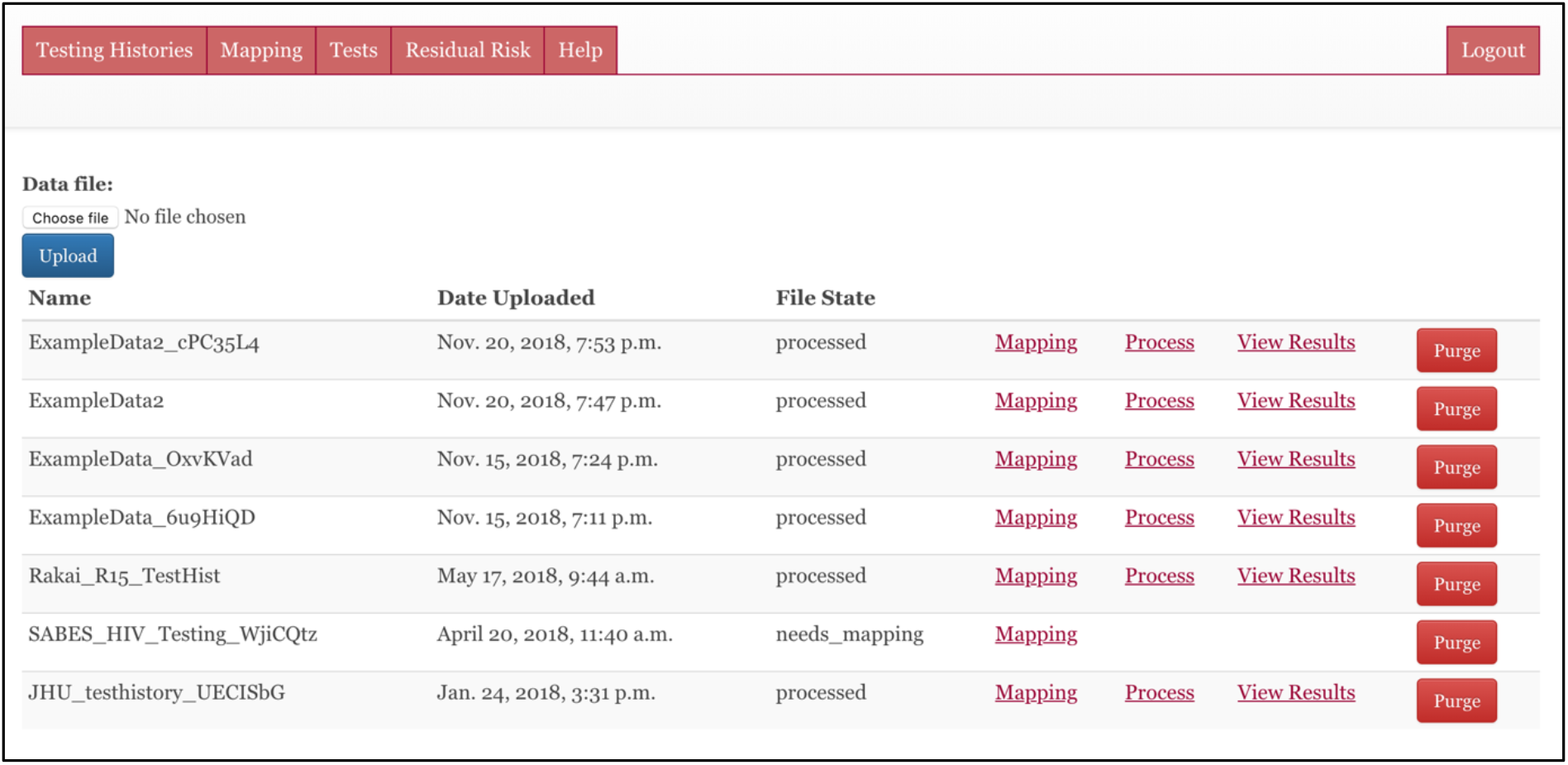
Testing Histories

### Mapping

This tab (Figure A.3) allows users to link strings (alphanumeric codes) in their data files to tests in the online database, hence linking records in uploaded files to the applicable diagnostic delays.

**Figure A.3:**
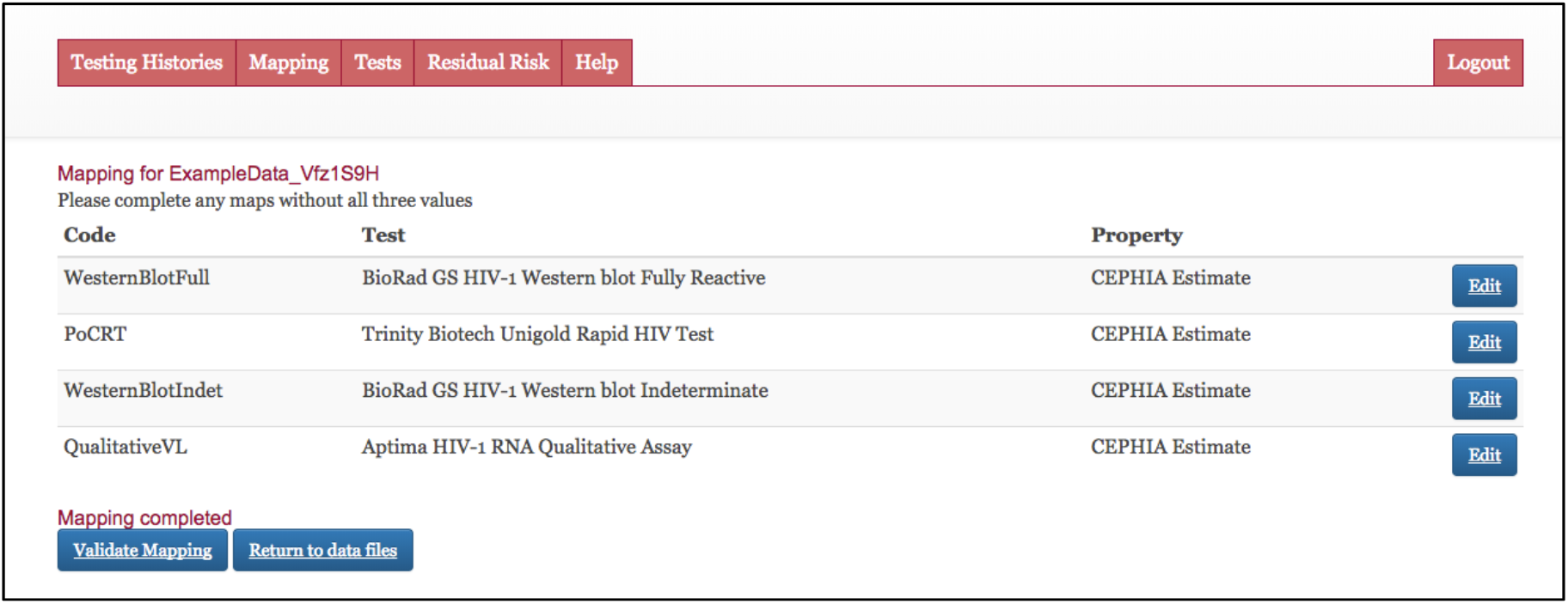
Mapping

### Tests

This tab (Figure A.4a) allows users to view the existing database of diagnostic tests, and to add new ones if necessary. Note that each user sees only the shared developer-maintained list of tests, plus his/her own – not those added by other users. This page further allows the user to select between computing EP-DDI and LP-DDI using naïve diagnostic delay medians, or to utilise the *σ* parameter and a specified value of *α* to compute credibility intervals (see Figures A.4b and A.4c).

**Figure A.4a:**
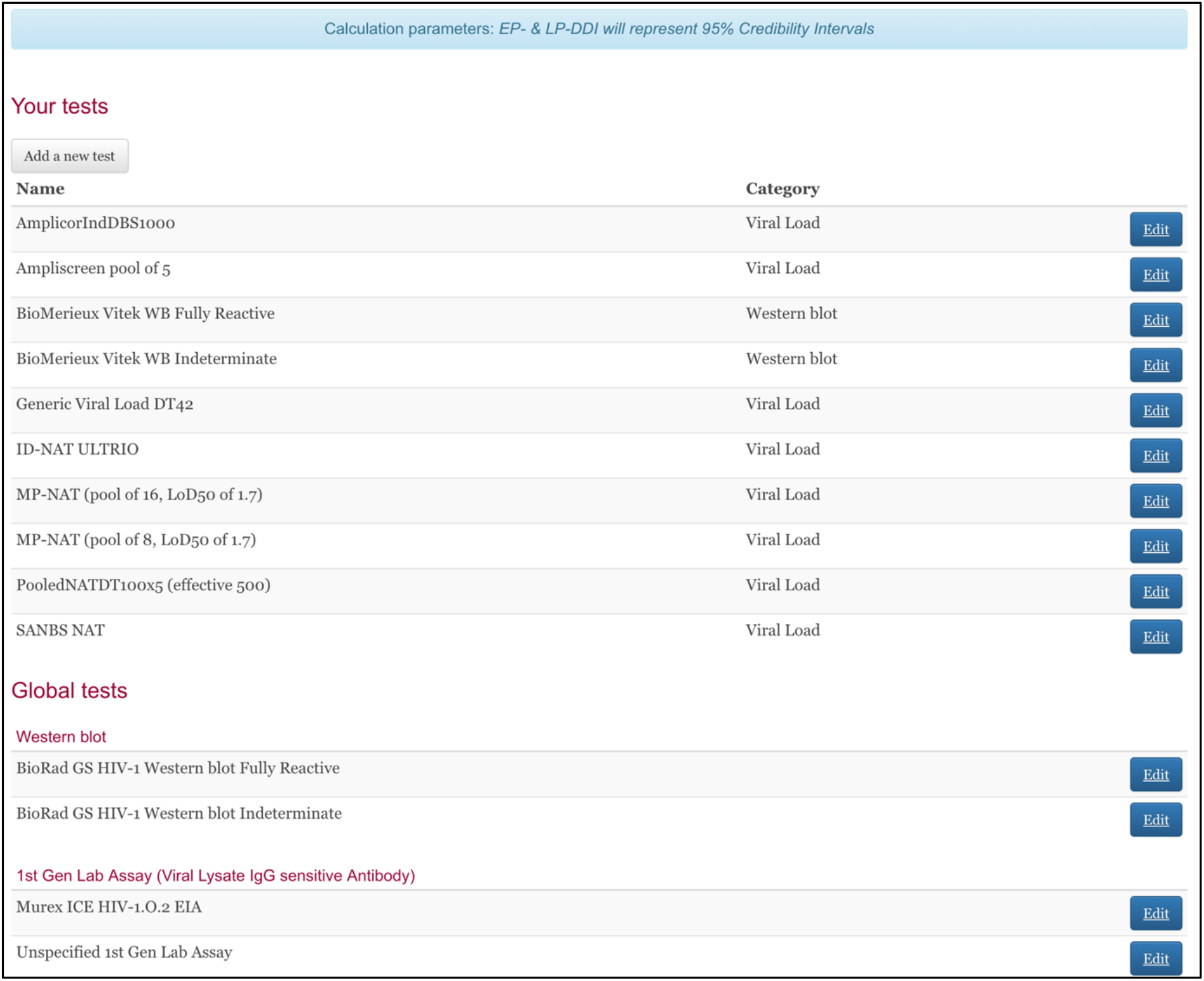
Tests

**Figure A.4b:**
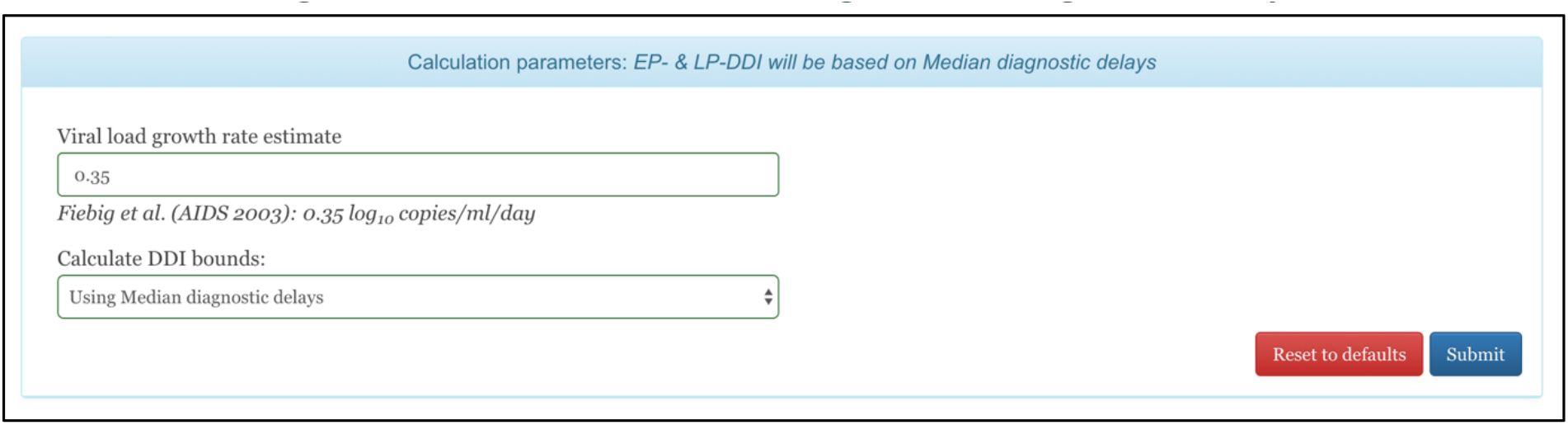
Naïve estimates using median diagnostic delays

**Figure A.4c:**
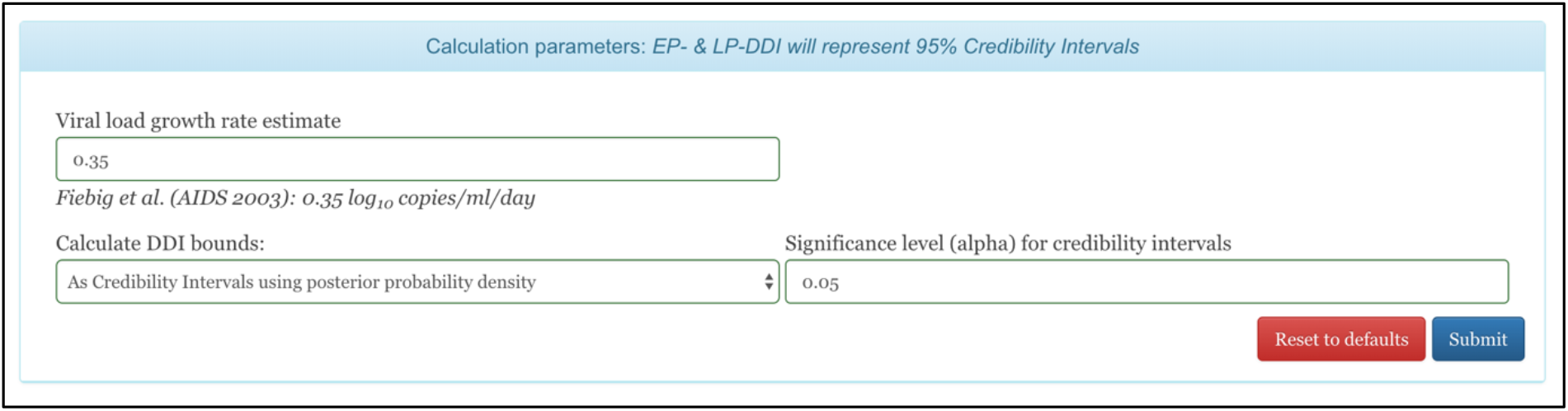
Computing credibility intervals

### Results

Processing can be triggered after test codes have been mapped to specific assays in the database. If test property estimates other than the default are preferred, these can be selected on the mapping screen prior to processing. Each file that has been uploaded on the “Testing Histories” tab has a “Mapping” link, and once mapping has been completed, a “Process” link appears. After processing, results can be viewed and downloaded on a per-file basis. Figure A.5a shows EP-DDI and LP-DDI based on median diagnostic delays, and Figure A.5b shows 95% credibility intervals.

**Figure A.5a:**
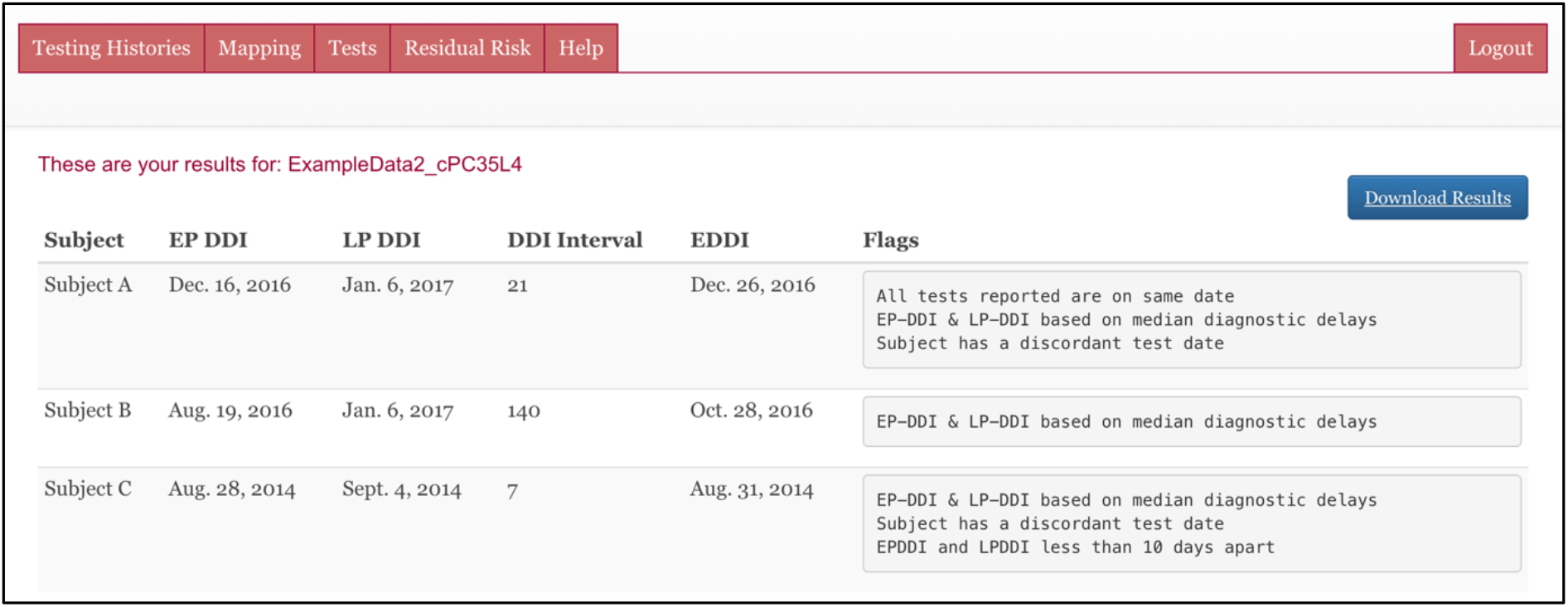
Results

**Figure A.5b:**
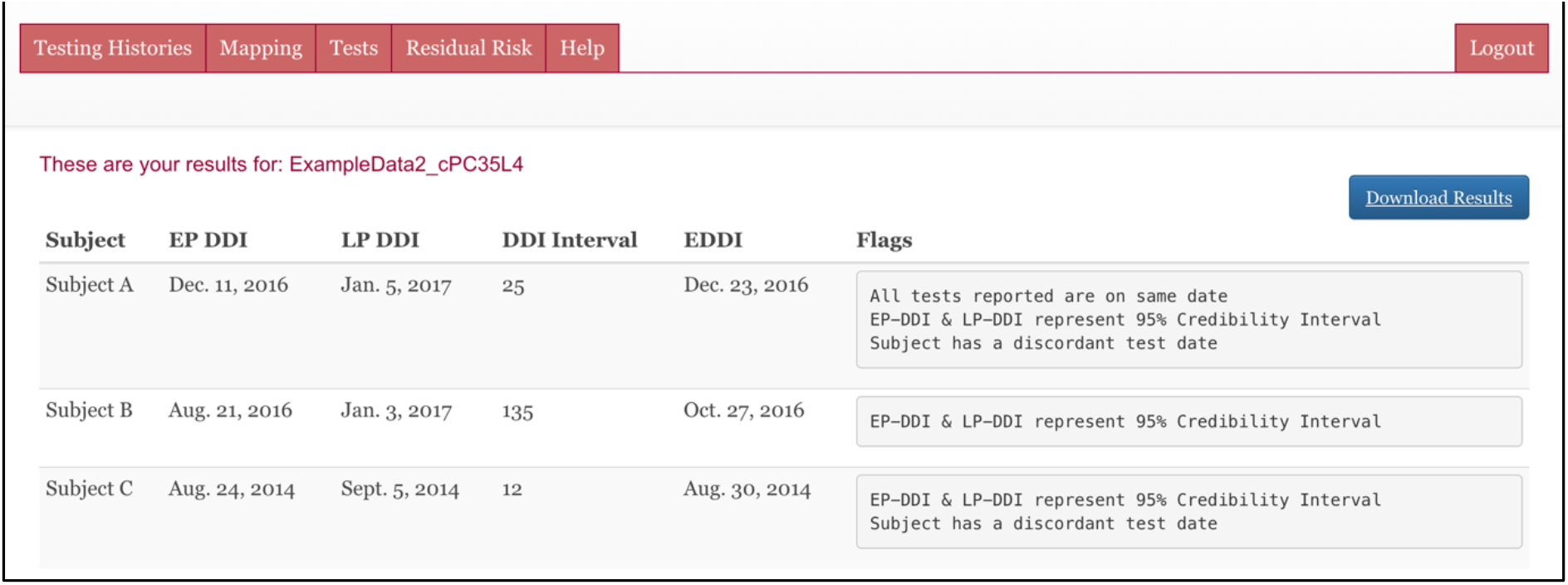
Results (95% CIs)

